# In Silico Investigations of Adaptive Therapy Using a Single Cytotoxic or a Single Cytostatic Drug

**DOI:** 10.1101/2023.05.14.540580

**Authors:** Daniel K. Saha, Alexander R. A. Anderson, Luis Cisneros, Carlo C. Maley

**Affiliations:** Arizona Cancer Evolution Center, Arizona State University; School of Life Sciences, Arizona State University; Biodesign Center for Biocomputing, Security and Society; Integrated Mathematical Oncology Department, Moffitt Cancer Center; Biodesign Center for Mechanisms of Evolution

## Abstract

Adaptive therapy, as per the dose modulation, dose-skipping, or intermittent treatment protocol works well for treatment using a single cytotoxic drug, under a wide range of scenarios and parameter settings. In contrast, adaptive therapy works well only under a limited number of scenarios and parameter settings when using a single cytostatic drug. In general, adaptive therapy works best under conditions of higher fitness cost, higher replacement rate, higher turnover. Adaptive therapy works best when drug dosages are changed as soon as a change in tumor burden is detected. In general, it is better to pause treatment sooner than later, when the tumor is shrinking If the amount of drug used is too low, it is unable to control the sensitive cells and the tumor grows. However, if the drug dose is too high, it quickly selects for resistant cells and eventually the tumor grows out of control. However, there appears to be intermediate levels of dosing, which we call the minimum effective dose, which is able to control the sensitive cells but is not high enough to select for the resistant cells to grow out of control.

## Introduction

Historically, treating cancer involves standard treatment (ST) at maximum tolerated dose (MTD) of a cancer drug. While this approach might work well for some cancer types, particularly ones with little heterogeneity, for most solid tumors this standard treatment eventually leads to an unresponsive tumor and consequent treatment failure instead of eradicating the cancer. This situation happens because under high doses of the drug resistant cell clones survive and proliferate at the expense of sensitive cells, a phenomenon termed ‘competitive release’ ^1^. Yet there is a silver lining: adaptations for resistance normally entail a fitness cost. For instance, resistant MCF7Dox cells have a doubling time of 60 hours versus 40 hours for sensitive MCF7 cells in the absence of the drug, and in co-culture experiments the sensitive MCF7 cells outcompete resistant MCF7Dox cells ^2^. Adaptive therapy utilizes this principle of fitness penalty incurred by resistant cells in absence of the drug in order to maintain long-term control over the tumor ^1–17^. It has been shown by Gatenby and colleagues that robust cancer control is possible with adaptive therapy as long as there is a substantial fitness cost to resistance. Multiple theoretical and mathematical models of adaptive therapy has been formulated. A vast array of models calibrate models to fit experimental data ^8, 9, 18^.

Preclinical experiments in mice with breast cancer demonstrated the superiority of dose modulation protocol, which involves adjusting drug dosages in response to changes in tumor burden over dose-skipping protocol, which involves administration of fixed-dosage of the drug if the tumor grows ^6^. Stable tumor burden was maintained in mice with breast cancer by a heuristic dose adjustment treatment protocol ^5^. Intermittent therapy trials, in which drug is administered until tumor burden shrinks to a fraction of the baseline followed by withholding drug until tumor burden increases to the baseline, in patients with prostate cancer resulted in an increase in median time to progression by at least 27 months compared to a contemporaneous cohort of patients ^7^. Preclinical adaptive therapy experiments were conducted in mice with breast cancer using paclitaxel ^5, 6^, or in mice with ovarian cancer using carboplatin ^5^. Both these drugs are cytotoxic in their mode of action. Previous adaptive therapy experiments have also been carried out using a cytotoxic drug ^6^. In contrast, the clinical trial carried out in prostate cancer patients using intermittent therapy used abiraterone (plus prednisone) ^7^, a hormone therapy drug with a cytostatic mode of action. The main distinction is that cytotoxic drugs work by killing cancer cells, while the mode of action of cytostatic drugs is the inhibition of tumor growth by suppressing cell division ^19^.

There are different variations and parameters for different types of adaptive therapy. While the dose-modulation protocol involves adjusting drug dosages based on changes in tumor burden since the previous treatment cycle, the dose-skipping or intermittent treatment protocols typically involve administering fixed dosages of the drugs every treatment cycle. Also, due to wide variations possible in cell kinetics, multiple scenarios exist. In addition to tumor growth kinetics, each adaptive therapy treatment protocol involves its own specific set of treatment parameters. Furthermore, because cytotoxic and cytostatic drugs vary fundamentally in their mode of action, survival outcomes could reasonably be expected to vary widely, thus designing adaptive therapy protocols that are best for a given situation and circumstance can be a challenging task. In this article, we sought to investigate which treatment protocol would be the best or most optimum for given situations or circumstances, on a case-by-case basis. We investigate three types of adaptive therapy treatment protocols, namely, dose modulation, dose-skipping, and intermittent therapy, as well as standard treatment at maximum tolerated dose under a wide range of conditions, with the goal of finding the best or the most optimum treatment protocol.

## Materials and Methods

We modified the agent-based hybrid model we previously published ^17^ to simulate adaptive therapy using a single drug and correspondingly extended the Hybrid Automata Library (HAL) agent-based modeling framework ^20^. Drug diffusion is modeled by solving the diffusion partial differential equations ^20^. The tumor consists of two different cell types: the sensitive cells and the resistant cells, which are situated on a 2-dimensional 100 by 100 square lattice. Each cell is modeled as an on-lattice agent, such that they occupy the lattice unit in which they are located and are not free to move around. Each cell, at every time step, could die or divide. As a first approximation, we assume that cytotoxic drugs only work by killing cells, and cytostatic drugs only work by inhibiting cell division. Thus, when cytotoxic drugs are used, sensitive cells die as a function of drug concentration as well as due to the background cell death rate, while resistant cells die only because of the background cell death rate. When cytostatic drugs are used, cells die only due to a background cell death rate (apoptosis rate) which is the same for all cell types. If a cell manages to survive, it could either divide or not, depending on the rate of cell division for that particular cell type. When a cell is committed to dividing, it could either divide so that a new cell occupies one of the available spaces in its Moore neighborhood, if any, or divide by replacing a neighbor, or do nothing if no empty space is available and cells are not allowed to replace a neighbor. Whether or not cells can replace a neighbor is governed by the replacement parameter. Setting the replacement parameter to zero enforces contact inhibition in the model and cells are not able to replace a neighbor while setting it to one allows dividing cells to replace a neighbor every time a cell is committed to dividing and an empty space is not available. Setting the replacement parameter to intermediate levels, such as 0.5, entails that cells committed to dividing would replace a neighbor 50% of the time, and not be able to replace a neighbor 50% of the time. When a cell divides, a daughter cell is created of the same type as the parent and is situated in one of the neighboring lattices. Both the daughter cells (the parent cell now having divided is considered one of the daughter cells) could mutate when cells divide. Whether or not the daughter cells would mutate after cell division is governed by a set of mutation parameters. We allow for bidirectional mutation in the model, meaning that a sensitive cell could either mutate to become a resistant cell, or not, as well as a resistant cell could mutate to become a sensitive cell, or not, at every time-step when cell division occurs. For the default parameter values, we set the mutation rate from sensitive to resistant cells, and vice-versa, to 10^-3^ per cell division, in order to account for the unrealistically low number of cells in our model. We have incorporated fitness cost in our model by choosing higher division rates for sensitive cells and lower division rates for resistant cells. Thus at any time step the division probability of sensitive cells > the division probability of resistant cells. When cytostatic drugs are being used, the sensitive cells undergo a decrease in their division rates as a function of the drug concentration, while the division rates for the resistant cells remain unaffected. At every time-step after the start of treatment drug diffusion is modeled by using the alternating direction implicit (ADI) method ^20^.

The treatment protocols being considered are the three different adaptive therapy protocols: dose modulation, dose-skipping, and intermittent; as well as standard treatment, which serves as the control. All adaptive therapy protocols involve monitoring the tumor every three days, and treatment starts as soon as the tumor burden equals or exceeds 50% of the carrying capacity, which is the total number of cancer cells that can be accommodated in the grid.

### Dose Modulation

The dose modulation protocols have two primary parameters: Delta Tumor, which is the percentage by which the tumor burden must change relative to the previous treatment cycle, in order to trigger a change in drug dose and, Delta Dose, which is the percentage by which the drug dose is changed relative to the previous treatment cycle. In this treatment the drug dosage is increased when the tumor has grown by more than Delta Tumor, or decreased when the tumor has shrunk by more than Delta Tumor, or maintained at the same dosage level as the previous treatment cycle when the tumor is either growing or shrinking by less than the Delta Tumor threshold. The drug dosage is capped so it never exceeds the maximum tolerated dose at which treatment is initiated, and never decreases beyond the minimum drug dose, which represents the amount below which the drug has no physiologic effect (and a dosage that in fact cannot be formulated in laboratory settings). In addition, if the absolute tumor burden ever exceeds the maximum of what has been recorded so far since initiation of treatment, drug dosage is increased by Delta Dose. Furthermore, a treatment vacation is triggered when the tumor burden falls to, or below a certain threshold (“stop dosing”), such that no drug is administered for that treatment cycle.

### Dose-Skipping

In contrast to the dose modulation protocol, the dose-skipping protocol involves administering a constant amount of the drug (fixed drug dosage). Drug is administered at that fixed level only when the tumor is growing above the Delta Tumor threshold, in all other cases no drug is administered (hence the treatment is “skipped” for that treatment cycle).

### Intermittent

The intermittent protocol involves monitoring the absolute tumor burden every treatment cycle. Treatment starts as soon as the tumor burden equals or exceeds 50% of the carrying capacity, which is considered to be 100% of the baseline level. A fixed-dose of the drug is administered at every treatment cycle until the tumor shrinks to 50% or more of the baseline level, at which point no drug is administered in any treatment cycle until the tumor burden increases to 100% of the baseline, and so on and so forth. The intermittent protocol has a key parameter: at what tumor burden should the treatment be stopped when the tumor is shrinking, in order that the tumor may be allowed to climb back up to the baseline value at which treatment was initiated previously. As mentioned above, this “stop dosing” threshold is chosen to be 50% of the baseline for the default parameter value.

The complete description of the model using the standard overview design details (ODD) format for describing agent-based models ^21^ can be found in ^17^ with the following changes:

In section 2.2 (Entities, State Variables, and Scales), we have considered two different cell types: sensitive, and resistant to account for treatment using either a single cytotoxic, or a single cytostatic drug.

In section 2.4.11 (Observation), we have made some modification to our criterion for progression. The modified survival criterion is as follows: If the tumor burden equaled or exceeded 97% of the carrying capacity at any point after initiation of therapy, or the rolling average of the total number of resistant cells over 500 time-steps equaled or exceeded 50% of the carrying capacity, then the particular run is scored as “Progressed” and the time at which progression takes place after therapy initiation is noted.

In section 2.5 (Initialization), instead of considering 4 different cell types to account for 2 drugs, in the initial tumor seed, we now consider 2 different cell types: sensitive and resistant to account for the cell types that are either sensitive, or resistant, to the single drug studied here.

In section 2.7.1 (Cell Death), for treatment with a single cytotoxic drug, the equation for probability of cell death is as follows: Probability of cell death per hour=background death probability per hour+S1*[Drug1]*Ψ1, where S1 is the binary indicator variable for the cell’s sensitivity to drug 1,[Drug1] being the concentration of drug 1 (non-negative real values), and Ψ1 is the drug potency (non-negative real values), quantified as the probability of cell death per unit drug concentration per hour. For treatment using a single cytostatic drug, the equation for probability of cell death is as follows: Probability of cell death per hour=background death probability per hour.

In section 2.7.2 (Cell Division), the cell division rates for the sensitive cell is 0.06 per hour, the cell division rate for the resistant cell is 0.02 per hour. The division probabilities can now be arranged in the following descending order:sensitive cells > resistant cells. For treatment with a single cytostatic drug, probability of cell division per hour=background division probability per hour-S1*[Drug1]*Ψ1, where S1 is the binary indicator variable for the cell’s sensitivity to drug 1, [Drug1] is the concentration of drug 1 (non-negative real values), and Ψ1 is the drug potency (non-negative real values), quantified as the probability of inhibition in cell division per unit drug concentration per hour.

Section 2.7.4 (Mutation): The default value for the mutation rate parameter is 10- 3 per cell division, to account for transition from sensitive cell type to resistant cell type, and vice-versa.

In section 2.7.6 (Drug Protocols): The treatment protocols are described as follows:

#### Standard Treatment (ST)

Drug 1 was administered at maximum tolerated dose (MTD) once every 24 hours for the entire duration of the simulation (Fig. 1).

**Figure 1:**
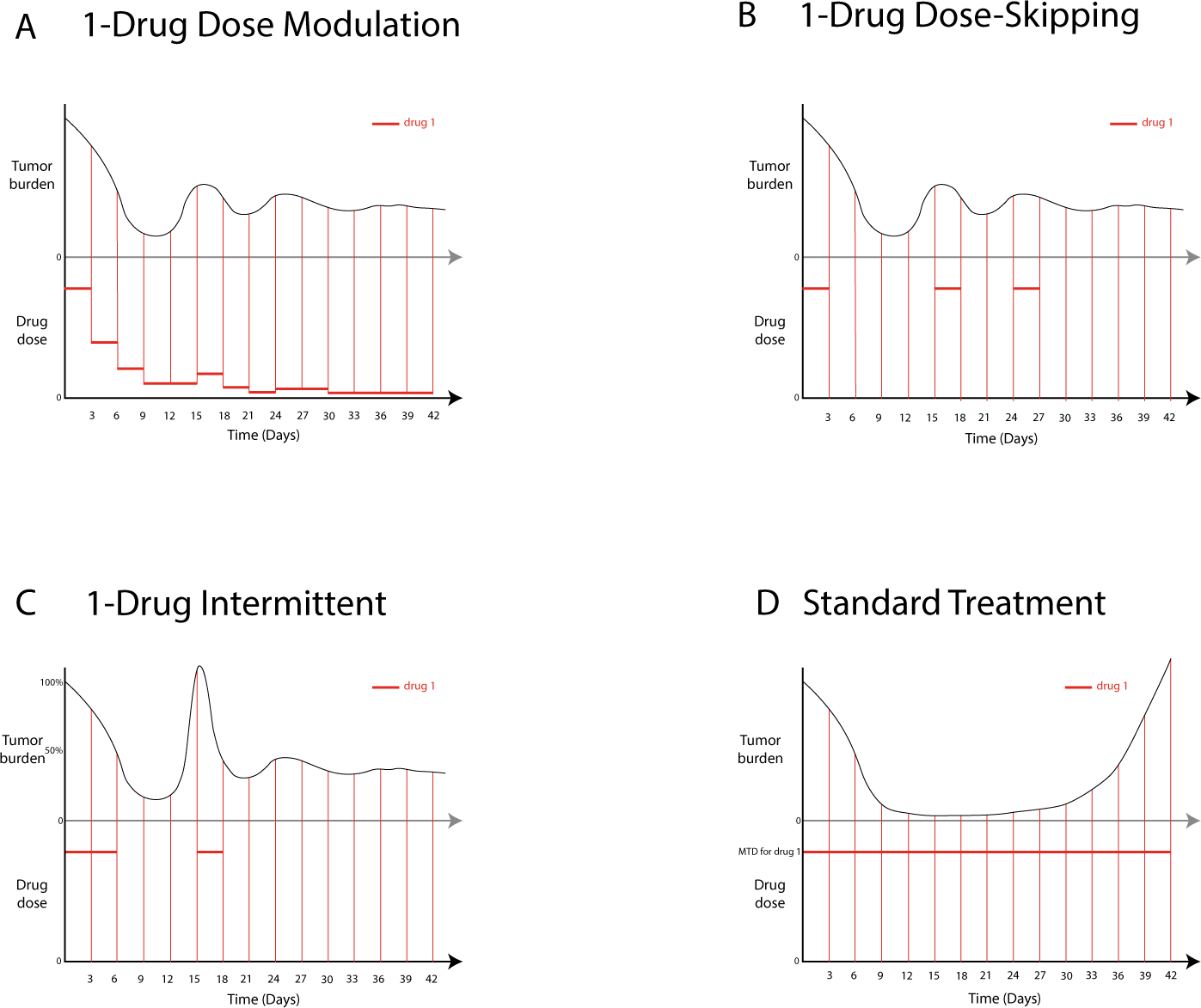
Single-drug adaptive therapy protocols using a single cytotoxic, or a single cytostatic drug. Schematic depicting dose modulation protocol (Fig. 1A), dose-skipping (Fig. 1B), intermittent (Fig. 1C), and standard treatment (Fig. 1D).

#### Dose Modulation

Treatment started at MTD with Drug 1, and dosage of the drug was adjusted according to the dose modulation adaptive therapy protocol, parameterized by Delta Tumor, and Delta Dose (Fig. 1). This treatment protocol was equivalent to the standard dose modulation adaptive therapy protocol (AT-1) from previous experiments.

#### Dose-Skipping

Drug 1 was administered at a fixed-dose that was set at 75% of MTD. If the tumor grew by more than Delta Tumor since its last measurement, or if the tumor burden exceeded its previous maximum size, the drug was applied or else, the dose was skipped. (Fig. 1). This is equivalent to AT-2 protocol from previous experiments.

#### Intermittent

Treatment started at 75% of the MTD using Drug 1, drug being administered once every 24 hours. Treatment was stopped when a shrinkage in tumor burden by at least 50% relative to the tumor burden at which therapy was initiated is detected, and therapy was restarted when the tumor burden equaled or exceeded 100% of the value at which therapy was initiated (Fig. 1). This protocol is equivalent to the prostate cancer clinical trial carried out in cancer patients with the drug abiraterone.

## Results

### Cytotoxic and Cytostatic Therapies

For treatment using a single cytotoxic drug, all the protocols, that is, dose modulation, dose-skipping, and intermittent work well, increasing TTP relative to standard treatment (Fig. 2A, Table 1), although the effect size as measured by the hazard ratio was small for intermittent (Table 1). For treatment using a single cytostatic drug, none of the protocols, that is, dose-modulation, intermittent, or dose-skipping is able to increase TTP relative to standard treatment (Fig. 2B, Table 1).

**Figure 2:**
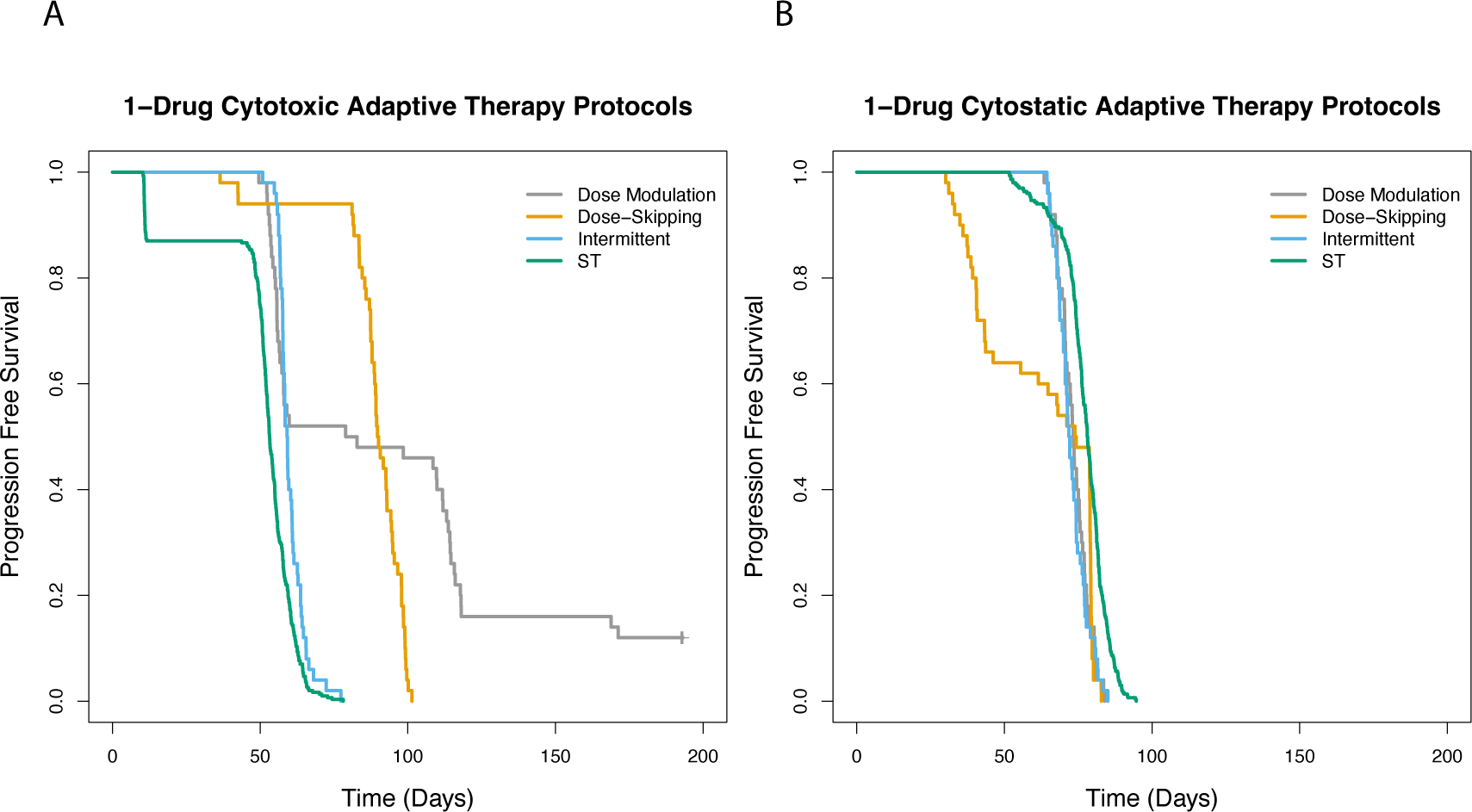
Adaptive therapy using a single cytotoxic or a single cytostatic drug. Single-drug adaptive therapy protocols comparing standard treatment (ST) versus three different adaptive therapy protocols, dose modulation, dose-skipping, and intermittent using a single cytotoxic drug (Fig. 2.2A), or a single cytostatic drug (Fig. 2.2B).

**Table 1:** Adaptive therapy using a single cytotoxic or a single cytostatic

| Experimental Condition | Comparison Condition | Hazard Ratio | 95% CI | *p*-value |
| --- | --- | --- | --- | --- |
| **Dose Modulation (cytotoxic)** |  |  |  |  |
| Default | Standard Treatment | 0.18 | 0.11-0.27 | <0.001 |
| **Dose Skipping (cytotoxic)** |  |  |  |  |
| Default | Standard Treatment | 0.01 | 0.003-0.037 | <0.001 |
| **Intermittent (cytotoxic)** |  |  |  |  |
| Default | Standard Treatment | 0.51 | 0.38-0.69 | <0.001 |
| **Dose Modulation (cytostatic)** |  |  |  |  |
| Default | Standard Treatment | 2.4 | 1.7-3.2 | <0.001 |
| **Dose Skipping (cytostatic)** |  |  |  |  |
| Default | Standard Treatment | 2.2 | 1.6-3.0 | <0.001 |
| **Intermittent (cytostatic)** |  |  |  |  |
| Default | Standard Treatment | 2.6 | 1.9-3.5 | <0.001 |

### The Effect of Fitness Costs of Resistance

Fitness cost incurred by resistant cells, as manifested in longer doubling times relative to sensitive cells in the absence of the drug, plays an important role in determining the outcome of both cytotoxic as well as cytostatic single-drug adaptive therapy. In general, adaptive therapy using a single cytotoxic drug works better than standard treatment at fitness cost of 1.7-fold, 2.5-fold, 5-fold, or 7-fold, increasing TTP, for treatment as per the dose modulation, intermittent, and dose-skipping protocol (Fig. 3, Table 2). The exception to this trend is adaptive therapy is not working well relative to the standard treatment at a fitness cost of 7-fold for treatment as per the dose-skipping protocol (Fig. 3C, Table 2). Moreover, in general, adaptive therapy treatment protocols under conditions of higher fitness cost leads to improved survival outcome relative to treatment under conditions of lower fitness cost. Thus, adaptive therapy at 7-fold fitness cost leads to better survival outcome than treatment under 5-fold fitness cost for dose-modulation as well as the intermittent treatment protocol (Table 2). However, for treatment with a cytostatic drug, none of the protocols tested here resulted in increased TTP relative to standard treatment at any of the fitness cost values tested here except dose-skipping protocol under fitness cost of 1.7-fold (Fig. 3B,3D,3F, Table 2).

**Figure 3:**
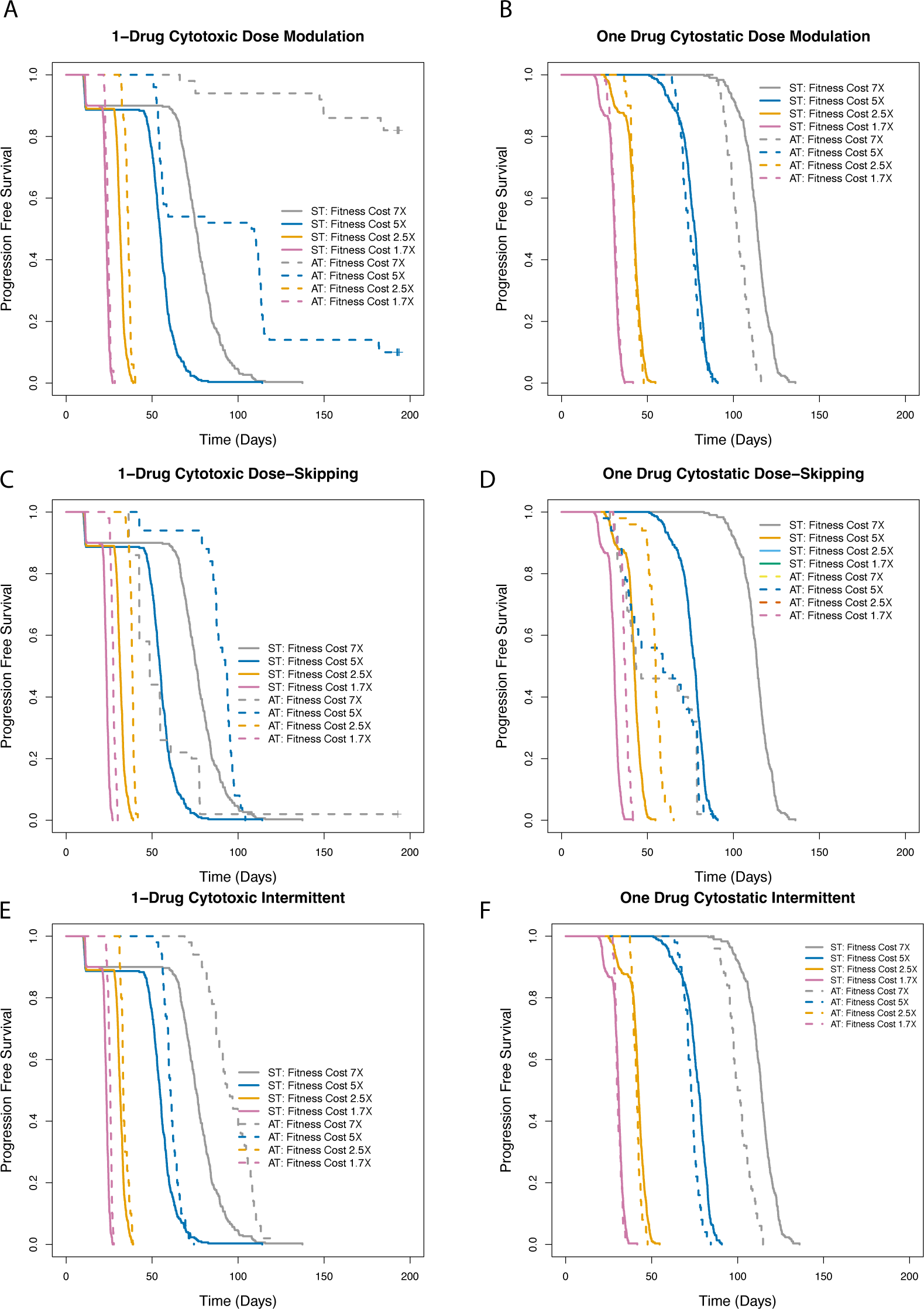
Effect of fitness cost parameter on the outcome of adaptive therapy using a single cytotoxic or a single cytostatic drug. Survival outcome for treatment as per the dose modulation protocol (Fig. 3A, Fig. 3B), dose-skipping (Fig. 3C, Fig. 3D), or intermittent (Fig. 3E, Fig. 3F) under fitness cost of 1.7-fold, 2.5-fold, 5-fold, or 7-fold relative to standard treatment for treatment using either a single cytotoxic (Fig. 3A, 3C, 3E), or a single cytostatic drug (Fig. 3B, 3D, 3F).

**Table 2:** Effect of fitness cost parameter on the outcome of adaptive therapy using a single cytotoxic or a single cytostatic drug

| **Experimental Condition** | **Comparison Condition** | **Hazard Ratio** | **95% CI** | ***p*-value** |
| --- | --- | --- | --- | --- |
| **Dose Modulation (cytotoxic)** |  |  |  |  |
| 1.7-fold fitness cost | Standard Treatment | 0.59 | 0.44–0.81 | < 0.001 |
| 2.5-fold fitness cost | Standard Treatment | 0.30 | 0.22-0.41 | <0.001 |
| 5-fold fitness cost | Standard Treatment | 0.21 | 0.14-0.31 | <0.001 |
| 7-fold fitness cost | Standard Treatment | 0.01 | 0.003-0.037 | <0.001 |
| 7-fold fitness cost | 5-fold fitness cost | 0.08 | 0.04-0.16 | <0.001 |
| **Dose Skipping (cytotoxic)** |  |  |  |  |
| 1.7-fold fitness cost | Standard Treatment | 0.08 | 0.05-0.13 | <0.001 |
| 2.5-fold fitness cost | Standard Treatment | 0.11 | 0.07-0.16 | <0.001 |
| 5-fold fitness cost | Standard Treatment | 0.11 | 0.07-0.16 | <0.001 |
| 7-fold fitness cost | Standard Treatment | 2.7 | 2.0-3.8 | <0.001 |
| 5-fold fitness cost | 7-fold fitness cost | 0.17 | 0.11-0.28 | <0.001 |
| **Intermittent (cytotoxic)** |  |  |  |  |
| 1.7-fold fitness cost | Standard Treatment | 0.31 | 0.22-0.42 | <0.001 |
| 2.5-fold fitness cost | Standard Treatment | 0.50 | 0.37-0.68 | <0.001 |
| 5-fold fitness cost | Standard Treatment | 0.55 | 0.41-0.74 | <0.001 |
| 7-fold fitness cost | Standard Treatment | 0.34 | 0.25-0.46 | <0.001 |
| 7-fold fitness cost | 5-fold fitness cost | 0.01 | 0.003-0.040 | <0.001 |
| **Dose Modulation (cytostatic)** |  |  |  |  |
| 1.7-fold fitness cost | Standard Treatment |  |  | Not Significant |
| 2.5-fold fitness cost | Standard Treatment |  |  | Not Significant |
| 5-fold fitness cost | Standard Treatment |  |  | Not Significant |
| 7-fold fitness cost | Standard Treatment | 4.9 | 3.5-6.8 | <0.001 |
| 2.5 -fold fitness cost | 1.7-fold fitness cost | 0.01 | 0.002-0.045 | <0.001 |
| **Dose Skipping (cytostatic)** |  |  |  |  |
| 1.7-fold fitness cost | Standard Treatment | 0.16 | 0.11-0.23 | <0.001 |
| 2.5-fold fitness cost | Standard Treatment | 0.06 | 0.03-0.10 | <0.001 |
| 5-fold fitness cost | Standard Treatment | 2.1 | 1.6-2.9 | <0.001 |
| 7-fold fitness cost | Standard Treatment |  |  | Not Significant |
| **Intermittent (cytostatic)** |  |  |  |  |
| 1.7-fold fitness cost | Standard Treatment |  |  | Not Significant |
| 2.5-fold fitness cost | Standard Treatment | 1.6 | 1.2-2.2 | <0.01 |
| 5-fold fitness cost | Standard Treatment | 2.2 | 1.6-3.0 | <0.001 |
| 7-fold fitness cost | Standard Treatment | 5.9 | 4.2-8.3 | <0.001 |

Moreover, in general, higher fitness cost (such as 7-fold fitness cost) translated to an improvement in survival outcome relative to lower fitness cost (such as 5-fold) (Table 2). Thus, for treatment using a single cytotoxic drug, as per the dose modulation protocol, as well as the intermittent resulted in increased TTP relative to standard treatment for all the fitness cost values tested (Fig. 3, Table 2, all *p* < 0.001). For treatment with the dose-skipping protocol using a cytotoxic drug (Fig. 3C; Table 2), all fitness values except the 7-fold fitness cost results in significant increase in TTP relative to standard treatment. For the intermittent protocol using a cytotoxic drug (Fig. 3E), all values of fitness cost tested here resulted in significantly increased TTP relative to standard treatment (Fig. 3A; Table 2). We observe increased TTP comparing higher fitness cost to lower fitness costs (Table 2). As an exception to this general trend, however, we observe increased TTP under 5- fold fitness cost compared to 7-fold fitness cost with dose-skipping using a cytotoxic drug.

### Cell Competition

The relationship between cell crowding, cell death and cell proliferation and direct cell competition is unknown. We encapsulated this complexity in a parameter (Cell Replacement) that specifies the likelihood that a cell can replace its neighbor if there are no empty spaces in its immediate neighborhood when it tries to divide.

In general, for treatment using a single cytotoxic drug, under conditions of 50% or 100% replacement rate, every adaptive therapy protocol works well, increasing TTP relative to standard treatment (Fig. 4, Table 3), an exception being treatment as per the dose-skipping protocol under conditions of 100% replacement (Fig. 4C, Table 3). However, under conditions of 0% replacement rate, no improvement in survival outcome was observed relative to standard treatment for any of the adaptive therapy protocols tested here. In general, adaptive therapy under conditions of higher replacement rates (more direct competition) results in improved survival outcome relative to treatment under conditions of lower replacement rates (Tables 3). Thus, survival outcome is better at 50% replacement rate relative to 0% replacement rate, or at 100% replacement relative to 50% for all of the adaptive therapy protocols using a single cytotoxic drug (Table 3), an exception being survival outcome is better under 50% relative to 100% replacement rate for treatment as per the dose-skipping protocol (Fig. 4, Table 3). In contrast, unlike treatment using a single cytotoxic drug, no improvement in survival outcome relative to standard treatment was observed when treated using a single cytostatic drug for any of the adaptive therapy protocols (Fig. 4B, 4D, 4F) tested here.

**Figure 4:**
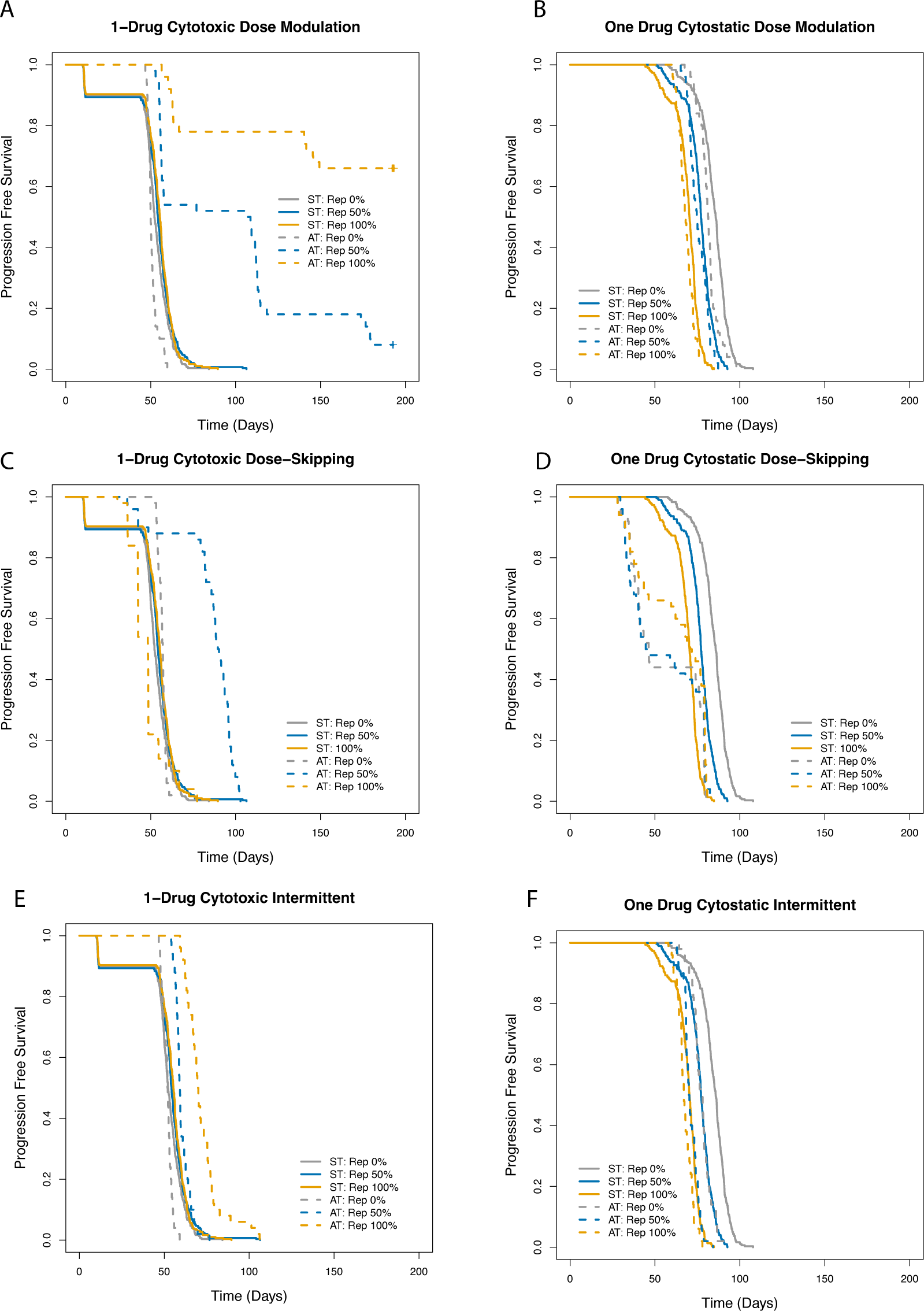
Effect of replacement parameter on outcome of adaptive therapy using a single cytotoxic or a single cytostatic drug. Treatment as per the dose modulation protocol (Fig. 2.4A, 2.4B), dose-skipping protocol (Fig. 4C, Fig. 4D), or intermittent (Fig. 4E, Fig. 4F), relative to standard treatment under conditions of 0%, 50%, or 100% replacement using either a single cytotoxic (Fig. 4A,4C, 4E), or a single cytostatic drug (Fig. 4B, Fig. 4D, Fig. 4F).

**Table 3.** : Effect of replacement parameter on outcome of adaptive therapy using a single cytotoxic or a single cytostatic drug

| **Experimental Condition** | **Comparison Condition** | **Hazard Ratio** | **95% CI** | ***p*-value** |
| --- | --- | --- | --- | --- |
| **Dose Modulation (cytotoxic)** |  |  |  |  |
| 0% replacement | Standard Treatment | 1.9 | 1.4-2.6 | <0.001 |
| 50% replacement | Standard Treatment | 0.17 | 0.11-0.27 | <0.001 |
| 100% replacement | Standard Treatment | 0.05 | 0.03-0.10 | <0.001 |
| 50% replacement | 0% replacement | 0.12 | 0.07-0.20 | <0.001 |
| 100% replacement | 50% replacement | 0.18 | 0.10-0.32 | <0.001 |
| **Dose Skipping (cytotoxic)** |  |  |  |  |
| 0% replacement | Standard Treatment |  |  | Not Significant |
| 50% replacement | Standard Treatment | 0.16 | 0.11-0.23 | <0.001 |
| 100% replacement | Standard Treatment | 1.7 | 1.3-2.3 | <0.001 |
| 50% replacement | 0% replacement | 0.04 | 0.02-0.11 | <0.001 |
| 100% replacement | 50% replacement | 24.8 | 10.0-61.4 | <0.001 |
| **Intermittent (cytotoxic)** |  |  |  |  |
| 0% replacement | Standard Treatment | 1.6 | 1.2-2.2 | <0.01 |
| 50% replacement | Standard Treatment | 0.60 | 0.45-0.81 | <0.001 |
| 100% replacement | Standard Treatment | 0.22 | 0.16-0.31 | <0.001 |
| 50% replacement | 0% replacement | 0.08 | 0.04-0.14 | <0.001 |
| 100% replacement | 50% replacement | 0.18 | 0.11-0.29 | <0.001 |
| **Dose Modulation (cytostatic)** |  |  |  |  |
| 0% replacement | Standard Treatment | 1.8 | 1.3-2.4 | <0.001 |
| 50% replacement | Standard Treatment | 1.4 | 1.0-1.9 | <0.05 |
| 100% replacement | Standard Treatment | 1.4 | 1.0-1.8 | <0.05 |
| **Dose Skipping (cytostatic)** |  |  |  |  |
| 0% replacement | Standard Treatment | 11.3 | 7.7-16.6 | <0.001 |
| 50% replacement | Standard Treatment | 2.4 | 1.8-3.3 | <0.001 |
| 100% replacement | Standard Treatment | 0.60 | 0.43-0.82 | <0.01 |
| **Intermittent (cytostatic)** |  |  |  |  |
| 0% replacement | Standard Treatment | 3.7 | 2.7-5.1 | <0.001 |
| 50% replacement | Standard Treatment | 3.4 | 2.5-4.7 | <0.001 |
| 100% replacement | Standard Treatment | 1.7 | 1.2-2.3 | <0.001 |

In general, higher replacement probabilities lead to better survival outcome relative to lower replacement probabilities. We observe increased TTP relative to standard treatment under conditions of higher replacement for treatment as per the dose modulation protocol using a cytotoxic drug (Table 3). An exception, however, to this general trend is treatment using a single cytotoxic drug using the dose-skipping protocol leads to increased TTP relative to standard treatment under conditions of 100% replacement versus 50% replacement (Fig. 4C, Table 3). In contrast, treatment with cytostatic drugs (Fig. 4B, 4D, 4F) do not result in increased TTP relative to standard treatment under any of the replacement conditions tested (Fig. 4B, 4D, 4F).

### Cell Turnover

For treatment with a single cytotoxic drug, under low turnover conditions, dose modulation (Fig. 5A) and intermittent treatment protocols (Fig. 5E) results in increased TTP relative to standard treatment but no increased TTP is observed for dose-skipping (Fig. 5C). For treatment with a single cytostatic drug, under conditions of low turnover, none of the protocols leads to increased TTP relative to standard treatment (Fig, 5B, 5D, 5F), standard treatment working well under these conditions. However, under conditions of high turnover, when treated with a single cytotoxic drug (Fig. 5A, 5C, 5E), or a single cytostatic drug (Fig. 5B, 5D, 5F), every adaptive therapy protocol tested here, that is, dose modulation (Fig. 5A, 5B), dose-skipping (Fig. 5C, 5D), and intermittent (Fig. 5E, 5F) treatment leads to increased TTP relative to standard treatment. In general, for treatment with a single cytotoxic drug, we observed improved survival outcomes under conditions of high turnover relative to conditions of low turnover (Table 4).

**Figure 5:**
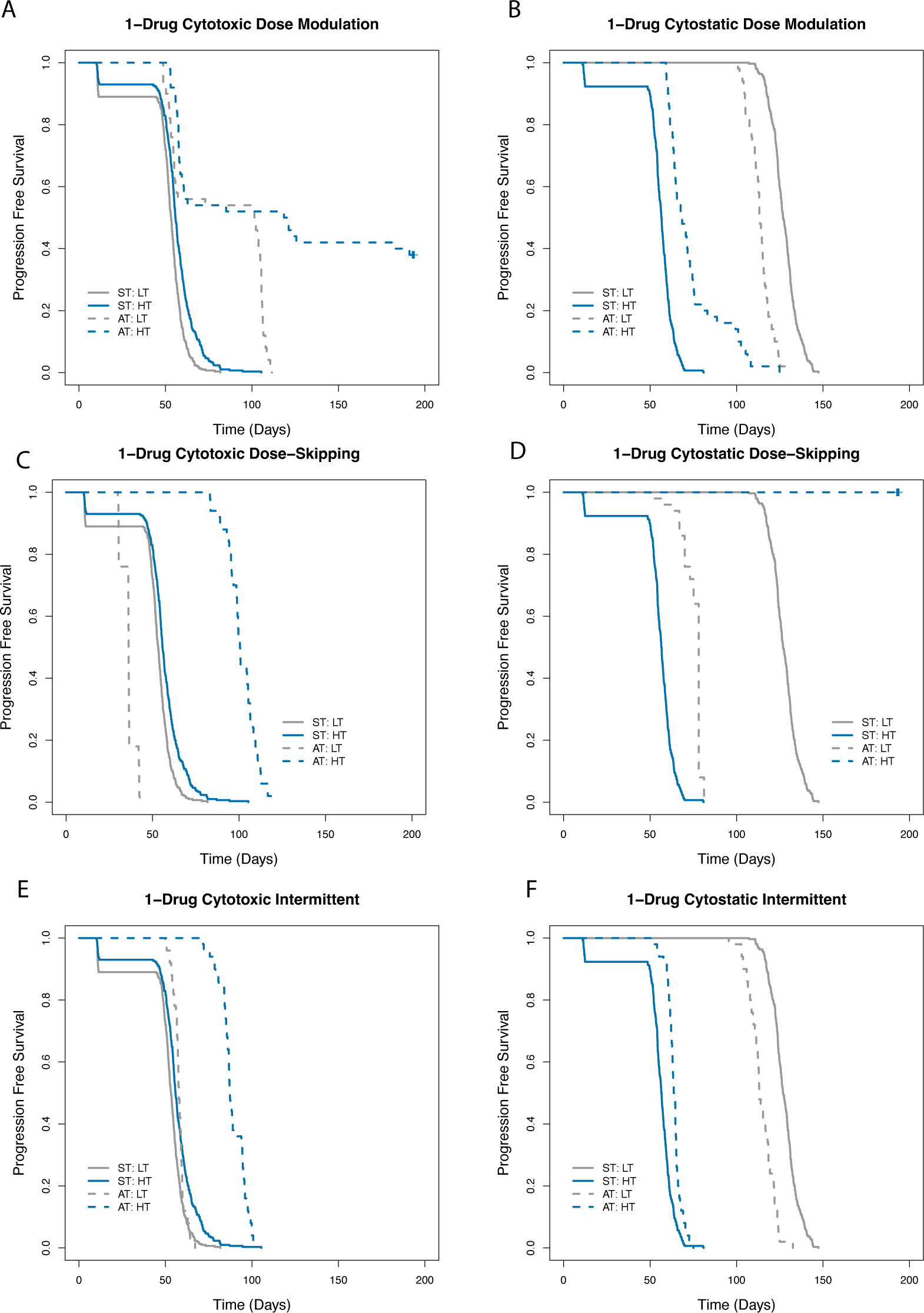
Effect of turnover on outcome of adaptive therapy using a single cytotoxic or a single cytostatic drug. Survival outcome for treatment as per the dose modulation protocol (Fig. 5A, Fig. 5B), dose-skipping protocol (Fig. 5C, Fig. 5D), or intermittent (Fig. 5E, Fig. 5F) using a single cytotoxic (Fig. 5A, 5C, 5E) or a single cytostatic drug (Fig. 5B, 5D, 5F) under conditions of low turnover (LT) or high turnover (HT), relative to standard treatment under those conditions.

**Table 4:** Effect of turnover on outcome of adaptive therapy using a single cytotoxic or a single cytostatic drug

| Experimental Condition | Comparison Condition | Hazard Ratio | 95% CI | *p*-value |
| --- | --- | --- | --- | --- |
| **Dose Modulation (cytotoxic)** |  |  |  |  |
| Low Turnover | Standard Treatment | 0.17 | 0.11-0.26 | <0.001 |
| High Turnover | Standard Treatment | 0.17 | 0.11-0.27 | <0.001 |
| High Turnover | Low Turnover | 0.30 | 0.18-0.50 | <0.001 |
| **Dose Skipping (cytotoxic)** |  |  |  |  |
| Low Turnover | Standard Treatment | 14.6 | 9.1-23.3 | <0.001 |
| High Turnover | Standard Treatment | 0.06 | 0.04-0.10 | <0.001 |
| High Turnover | Low Turnover | ~0 |  |  |
| **Intermittent (cytotoxic)** |  |  |  |  |
| Low Turnover | Standard Treatment | 0.63 | 0.47-0.85 | <0.01 |
| High Turnover | Standard Treatment | 0.13 | 0.09-0.19 | <0.001 |
| High Turnover | Low Turnover | ~0 |  |  |
| **Dose Modulation (cytostatic)** |  |  |  |  |
| Low Turnover | Standard Treatment | 8.5 | 6.0-11.9 | <0.001 |
| High Turnover | Standard Treatment | 0.19 | 0.13-0.27 | <0.001 |
| High Turnover | Low Turnover | 6.4 | 4.1-10.1 | <0.001 |
| **Dose Skipping (cytostatic)** |  |  |  |  |
| Low Turnover | Standard Treatment |  |  | Not Significant |
| High Turnover | Standard Treatment | ~0 |  |  |
| High Turnover | Low Turnover | ~0 |  |  |
| **Intermittent (cytostatic)** |  |  |  |  |
| Low Turnover | Standard Treatment | 6.7 | 4.8-9.3 | <0.001 |
| High Turnover | Standard Treatment | 0.40 | 0.29-0.54 | <0.001 |
| Low Turnover | High Turnover | $\sim 0$ | | |

### When to Adjust the Dose of the Drug

The dose modulation protocols have two primary parameters: Delta Tumor, which is the amount the tumor burden must change in order to trigger a change in drug dose and, Delta Dose, which is the amount by which the dose is changed.

For treatment using a single cytotoxic drug, as per the dose-modulation protocol, delta tumor=5%, or delta tumor=10% leads to increased TTP relative to standard treatment (Fig. 6A, Table 5), while treatment as per the dose-skipping protocol works well for all values of delta tumor tested here, that is, delta tumor=5%, 10%, 20%, or 40% (Fig. 6C, Table 5). However, only treatment as per the dose modulation protocol with delta tumor=5% increased TTP relative to standard treatment (Fig. 6B, Table 5). In general, the lower the value of delta tumor, the better is the survival outcome. Thus, delta tumor=5% leads to better survival outcome relative to delta tumor=10% for treatment as per the dose modulation protocol when using a single cytotoxic, or a single cytostatic drug (Table 5). These results indicate that treatment as per the dose modulation protocol works best if dose is adjusted as soon as a change in tumor burden is detected. In practice this will likely be limited by the sensitivity of the tumor burden assay. Note that using a small Delta Tumor value allows the dose modulation protocol to be effective even with a cytostatic drug (Fig. 6B, Table 5).

**Figure 6:**
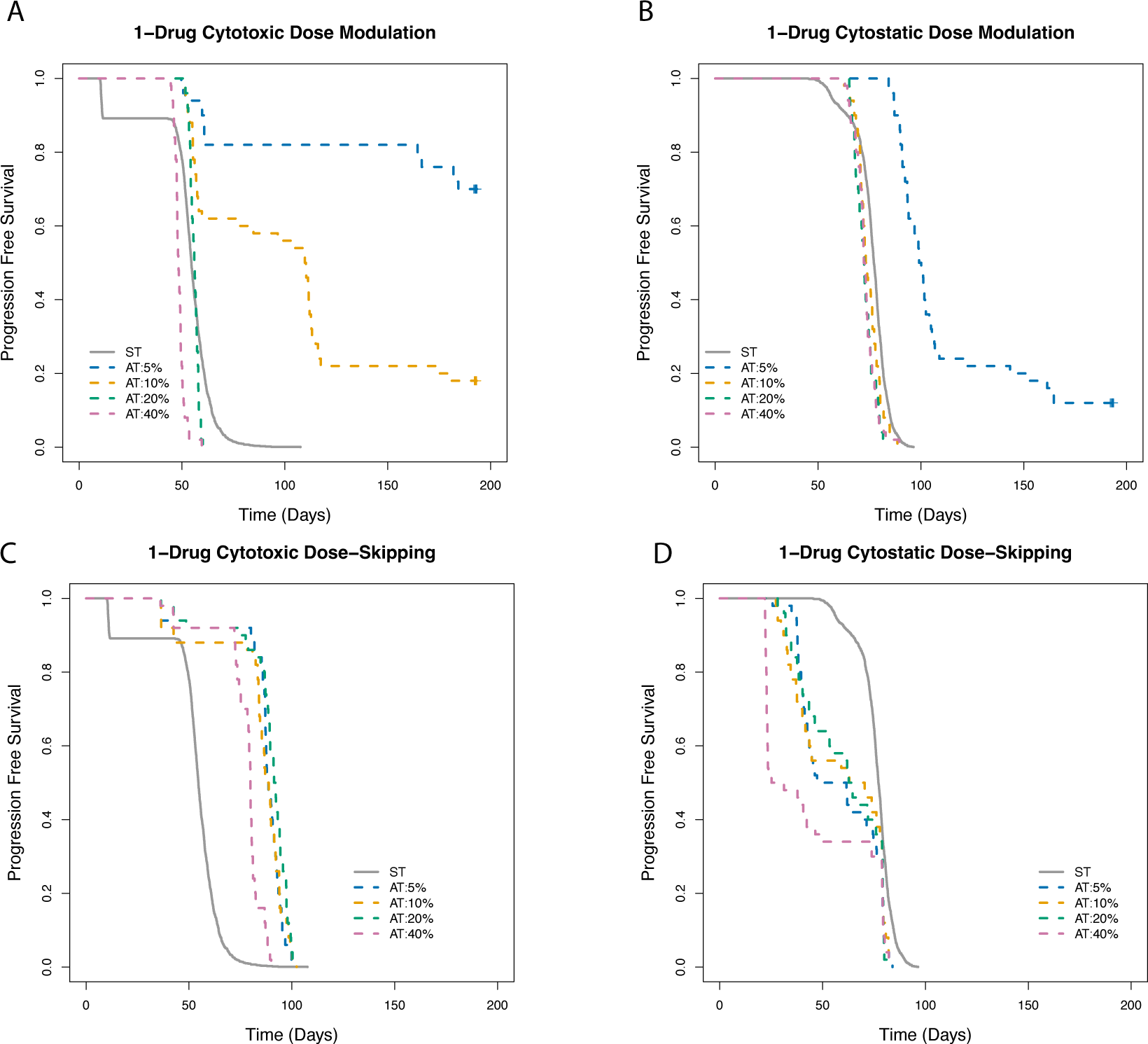
Effect of the delta tumor parameter on determining the outcome of adaptive therapy using a single cytotoxic or a single cytostatic drug. Survival outcome comparing dose modulation treatment protocol with Delta Tumor=5%, 10%, 20%, or 40% using a single cytotoxic (Fig. 6A), or a single cytostatic drug (Fig. 6B) relative to standard treatment. Survival outcome comparing dose-skipping treatment protocol with Delta Tumor=5%, 10%, 20%, or 40% using a single cytotoxic (Fig. 6C), or a single cytostatic drug (Fig. 6D).

**Table 5.** : Effect of the delta tumor parameter on determining the outcome of adaptive therapy using a single cytotoxic or a single cytostatic drug

| Experimental Condition | Comparison Condition | Hazard Ratio | 95% CI | *p*-value |
| --- | --- | --- | --- | --- |
| **Dose Modulation (cytotoxic)** |  |  |  |  |
| Delta Tumor=5% | Standard Treatment | 0.03 | 0.02-0.06 | <0.001 |
| Delta Tumor=10% | Standard Treatment | 0.11 | 0.07-0.17 | <0.001 |
| Delta Tumor=20% | Standard Treatment |  |  | Not Significant |
| Delta Tumor=40% | Standard Treatment | 4.4 | 3.3-6.0 | <0.001 |
| Delta Tumor=5% | Delta Tumor-10% | 0.21 | 0.11-0.38 | <0.001 |
| **Dose Skipping (cytotoxic)** |  |  |  |  |
| Delta Tumor=5% | Standard Treatment | 0.12 | 0.09-0.17 | <0.001 |
| Delta Tumor=10% | Standard Treatment | 0.13 | 0.10-0.19 | <0.001 |
| Delta Tumor=20% | Standard Treatment | 0.11 | 0.08-0.16 | <0.001 |
| Delta Tumor=40% | Standard Treatment | 0.19 | 0.14-0.26 | <0.001 |
| **Dose Modulation (cytostatic)** |  |  |  |  |
| Delta Tumor=5% | Standard Treatment | 0.06 | 0.04-0.10 | <0.001 |
| Delta Tumor=10% | Standard Treatment | 1.5 | 1.1-2.0 | <0.01 |
| Delta Tumor=20% | Standard Treatment | 2.3 | 1.8-3.1 | <0.001 |
| Delta Tumor=40% | Standard Treatment | 2.1 | 1.6-2.8 | <0.001 |
| Delta Tumor=5% | Delta Tumor=10% | 0.02 | 0.006-0.049 | <0.001 |
| **Dose Skipping (cytostatic)** |  |  |  |  |
| Delta Tumor=5% | Standard Treatment | 2.5 | 1.8-3.3 | <0.001 |
| Delta Tumor=10% | Standard Treatment | 2.0 | 1.5-2.6 | <0.001 |
| Delta Tumor=20% | Standard Treatment | 2.3 | 1.7-3.0 | <0.001 |
| Delta Tumor=40% | Standard Treatment | 2.7 | 2.1-3.6 | <0.001 |

### How much to change the dose for the dose modulation protocols

The dose modulation protocols have two primary parameters: Delta Tumor, which is the amount the tumor burden must change in order to trigger a change of drug dose and, Delta Dose, which is the amount by which the drug dose is changed.

For treatment using a single cytotoxic drug, as per the dose modulation protocol, all delta dose values tested here, that is, delta dose=25%, 50%, or 75% increased TTP relative to standard treatment (Fig. 7, Table 6). However, when using a single cytostatic drug, as per the dose modulation protocol, only delta dose=75% leads to increased TTP relative to standard treatment (Fig. 7, Table 6). In general, choosing a high value of delta dose improves survival outcome relative to choosing a lower value for delta dose (Fig. 7, Table 6). Thus, using delta dose=50% leads to better survival outcome than using delta dose=25% for treatment using a single cytotoxic drug, and using delta dose=75% leads to better survival outcome than using delta dose=50% when using a single cytostatic drug (Table 6).

**Figure 7:**
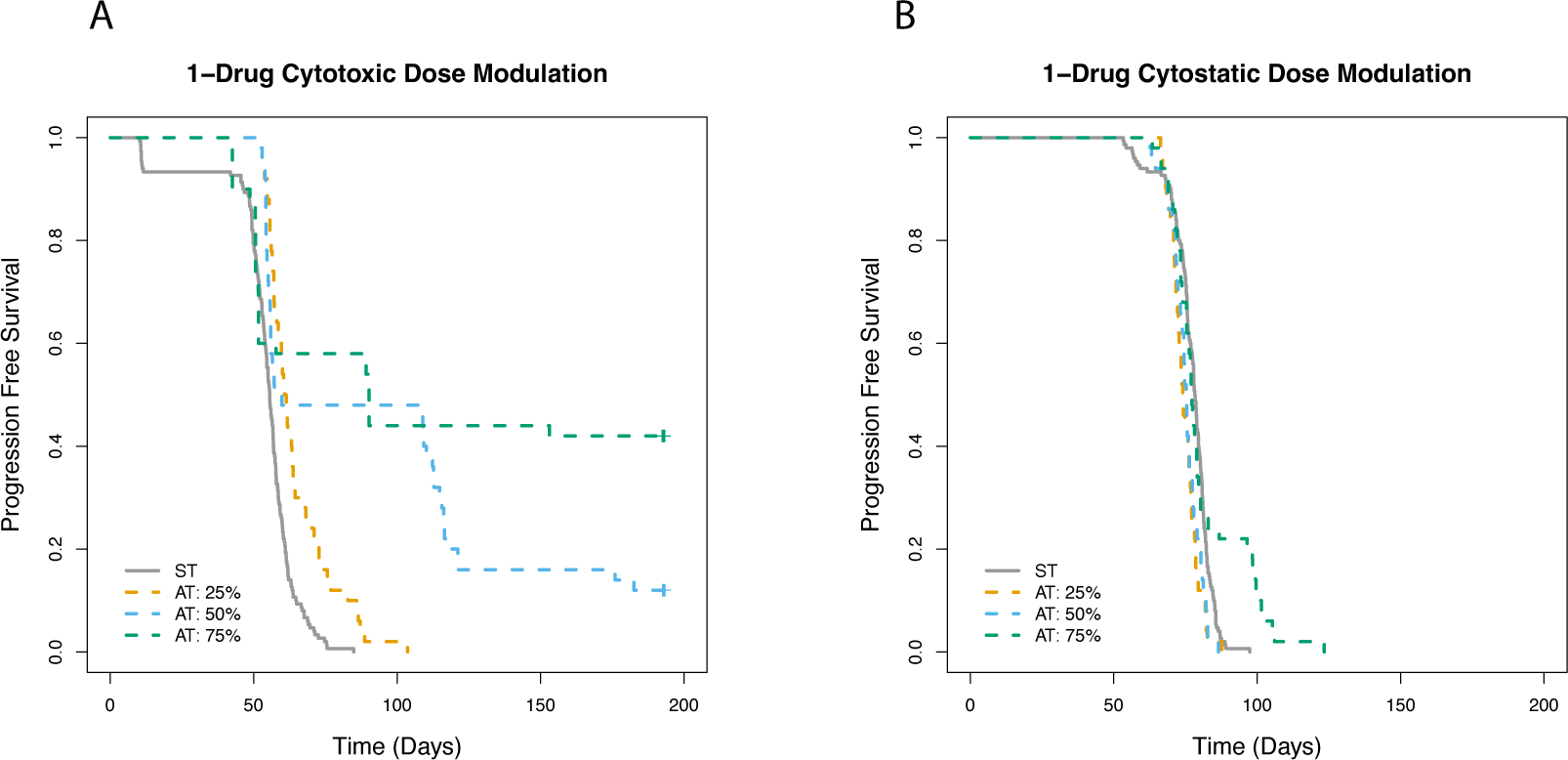
Effect of the delta dose parameter on determining the outcome of dose modulation adaptive therapy using a single cytotoxic or a single cytostatic drug. Survival outcome for treatment as per the dose modulation protocol with Delta Dose=25%, 50%, or 75% relative to standard treatment using a single cytotoxic drug (Fig. 7A), or a single cytostatic drug (Fig. 7B).

**Table 6:** Effect of the delta dose parameter on determining the outcome of dose modulation adaptive therapy using a single cytotoxic or a single cytostatic drug

| Experimental Condition | Comparison Condition | Hazard Ratio | 95% CI | p-value |
| --- | --- | --- | --- | --- |
| **Dose Modulation (cytotoxic)** |  |  |  |  |
| Delta Dose=25% | Standard Treatment | 0.41 | 0.29-0.58 | <0.001 |
| Delta Dose=50% | Standard Treatment | 0.26 | 0.16-0.40 | <0.001 |
| Delta Dose=75% | Standard Treatment | 0.20 | 0.12-0.33 | <0.001 |
| Delta Dose=50% | Delta Dose=25% | 0.43 | 0.26-0.71 | <0.001 |
| **Dose Modulation (cytostatic)** |  |  |  |  |
| Delta Dose=25% | Standard Treatment | 1.9 | 1.4-2.6 | <0.001 |
| Delta Dose=50% | Standard Treatment | 1.7 | 1.2-2.4 | <0.01 |
| Delta Dose=75% | Standard Treatment | 0.65 | 0.45-0.94 | <0.05 |
| Delta<br>Dose=75% | Delta<br>Dose=50% | 0.54 | 0.35-0.83 | <0.01 |

### Effects of stopping treatment when tumor burden falls below a certain level

The intermittent protocol has a key parameter: at what tumor burden should the treatment be stopped when the tumor is shrinking, in order that the tumor may be allowed to climb back up to the baseline value at which treatment was initiated previously. For treatment using a single cytotoxic drug, as per the intermittent protocol, pausing treatment when tumor shrinks by 20%, or 50% leads to increased TTP relative to standard treatment, but no improvement in survival outcome was observed relative to standard treatment when pausing treatment when tumor shrinks by 90%. However, no improvement in survival outcome was observed for any of the values tested here when using a single cytostatic drug (Fig. 8, Table 7). In general, the sooner treatment is paused the better is the survival outcome (Table 7).

**Figure 8:**
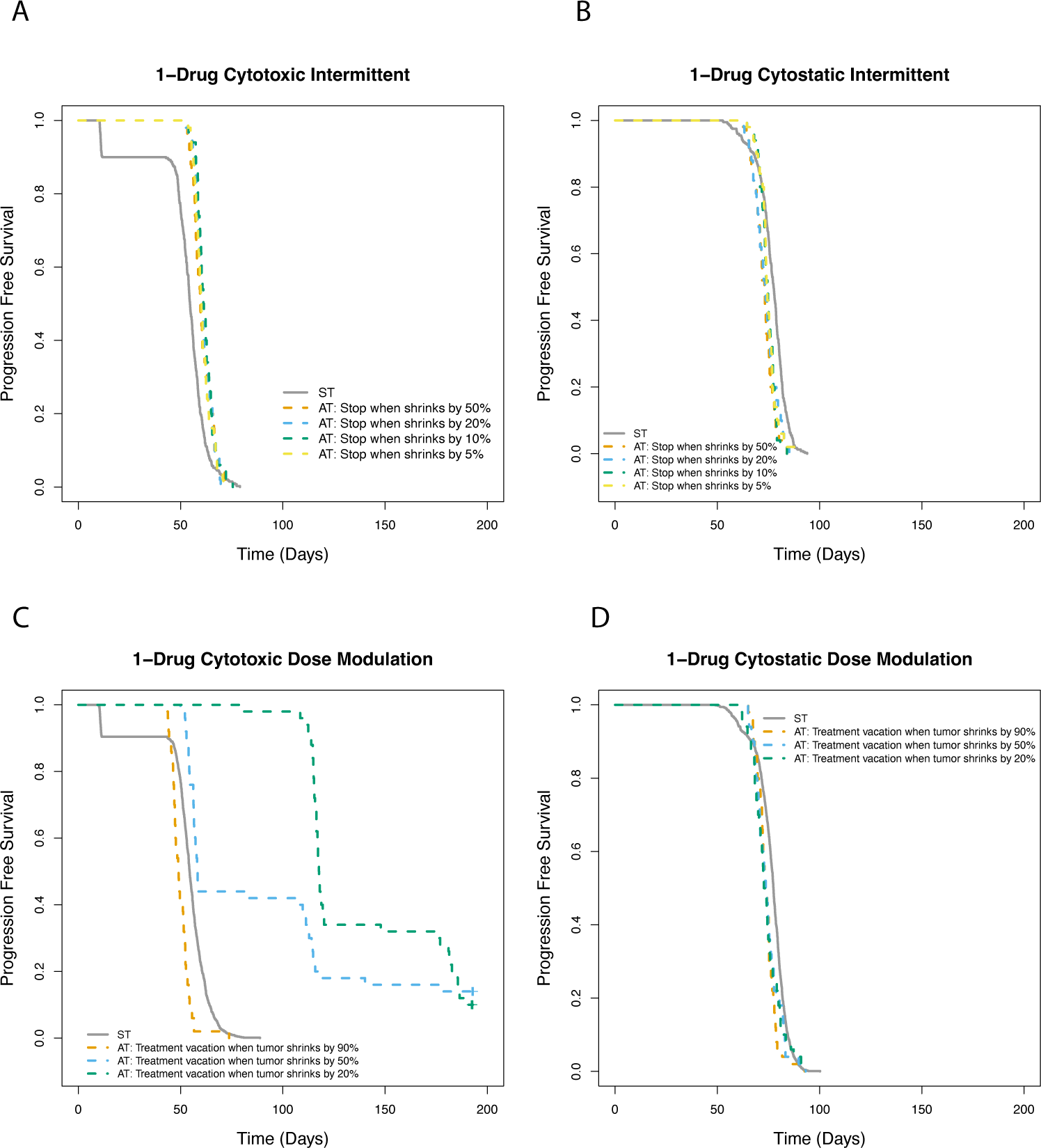
Effect of stopping treatment when tumor burden falls below a certain level for adaptive therapy using a single cytotoxic or a single cytostatic drug. For treatment as per the intermittent protocol using either a single cytotoxic (Fig. 8A), or a single cytostatic drug (Fig. 8B), the threshold for stopping treatment was varied as the tumor shrinks by 5%, 10%, 20%, or 50% of the pre-treatment baseline. Survival outcome for treatment using a single cytotoxic drug (Fig. 8C), or a single cytostatic drug (Fig. 8D) as the trigger for treatment vacation is when the tumor shrinks by 20%, 50%, or 90%.

**Table 7:** Effect of stopping treatment when tumor burden falls below a certain level for adaptive therapy using a single cytotoxic or a single cytostatic drug

| Experimental Condition | Comparison Condition | Hazard Ratio | 95% CI | *p*-value |
| --- | --- | --- | --- | --- |
| **Dose Modulation (cytotoxic)** |  |  |  |  |
| Treatment vacation when tumor shrinks by 20% | Standard Treatment | 0.003 | 0.0004-0.0191 | <0.001 |
| Treatment vacation when tumor shrinks by 50% | Standard Treatment | 0.20 | 0.13-0.30 | <0.001 |
| Treatment vacation when tumor shrinks by 90% | Standard treatment | 2.5 | 1.9-3.4 | <0.001 |
| Treatment vacation when tumor shrinks by 20% | Treatment vacation when tumor shrinks by 50% | 0.50 | 0.33-0.77 | <0.01 |
| Treatment vacation when tumor shrinks by 50% | Treatment vacation when tumor shrinks by 90% | 0.12 | 0.07-0.20 | <0.001 |
| **Intermittent (cytotoxic)** |  |  |  |  |
| Stop when shrinks by 5% | Standard Treatment | 0.54 | 0.40-0.73 | <0.001 |
| Stop when shrinks by 10% | Standard Treatment | 0.47 | 0.35-0.64 | <0.001 |
| Stop when shrinks by 20% | Standard Treatment | 0.51 | 0.38-0.69 | <0.001 |
| Stop when shrinks by 50% | Standard Treatment | 0.55 | 0.41-0.73 | <0.001 |
| **Dose Modulation (cytostatic)** |  |  |  |  |
| Treatment vacation when tumor shrinks by 20% | Standard Treatment | 1.4 | 1.1-1.9 | <0.05 |
| Treatment vacation when tumor shrinks by 50% | Standard Treatment | 1.4 | 1.0-1.8 | <0.05 |
| Treatment vacation when tumor shrinks by 90% | Standard treatment | 1.8 | 1.4-2.4 | <0.001 |
| **Intermittent (cytostatic)** |  |  |  |  |
| Stop when shrinks by 5% | Standard Treatment | 1.8 | 1.3-2.4 | <0.001 |
| Stop when shrinks by 10% | Standard Treatment | 2.1 | 1.5-2.8 | <0.001 |
| Stop when shrinks by 20% | Standard Treatment | 1.9 | 1.4-2.6 | <0.001 |
| Stop when shrinks by 50% | Standard Treatment | 2.5 | 1.8-3.4 | <0.001 |

For dose modulation protocols, an important consideration to be made is whether treatment vacation should be triggered when tumor shrinks by at least a certain percentage relative to the start of therapy. For treatment using a single cytotoxic drug, as per the dose modulation protocol, triggering a treatment vacation when the tumor shrinks by 20%, or 50% increases TTP relative to standard treatment, but no increase in TTP relative to standard treatment is observed when a treatment vacation is triggered when the tumor shrinks by 90% (Fig. 8, Table 7). In general, the sooner a treatment vacation is triggered when the tumor is shrinking the better is the survival outcome (Table 7). Thus, triggering a treatment vacation when the tumor shrinks by 20% results in improved survival outcome than triggering a treatment vacation when the tumor shrinks by 50% (Table 7), and triggering a treatment vacation when the tumor shrinks by 50% leads to a better survival outcome than triggering a treatment vacation when the tumor shrinks by 90% (Table 7). When using a single cytostatic drug, there was no improvement in survival outcome for any of these values tested (Fig. 8D).

### Drug dosage level at which adaptive therapy is initiated and capped

For the dose modulation protocol, we tested different drug dosage levels for initiating treatment. We also capped the dose level at that value, so that dose modulation was never allowed to exceed that level. We observe that for treatment using a single cytotoxic drug, as per the dose modulation protocol, initiating treatment at 50%, 75%, or 100% of MTD resulted in an increase in TTP relative to standard treatment (Fig. 9A, Table 8), whereas for treatment initiation at 25% of MTD, we observed no cases of death for either the standard treatment or treatment as per the dose modulation protocol (Fig. 9A), and thus there was no increase in TTP with the dose modulation protocol relative to standard treatment. For treatment using a single cytostatic drug, as per the dose modulation protocol, none of the values tested, that is, 25%, 50%, 75%, or 100% of MTD resulted in increase in TTP relative to standard treatment (Fig. 9B).

**Figure 9:**
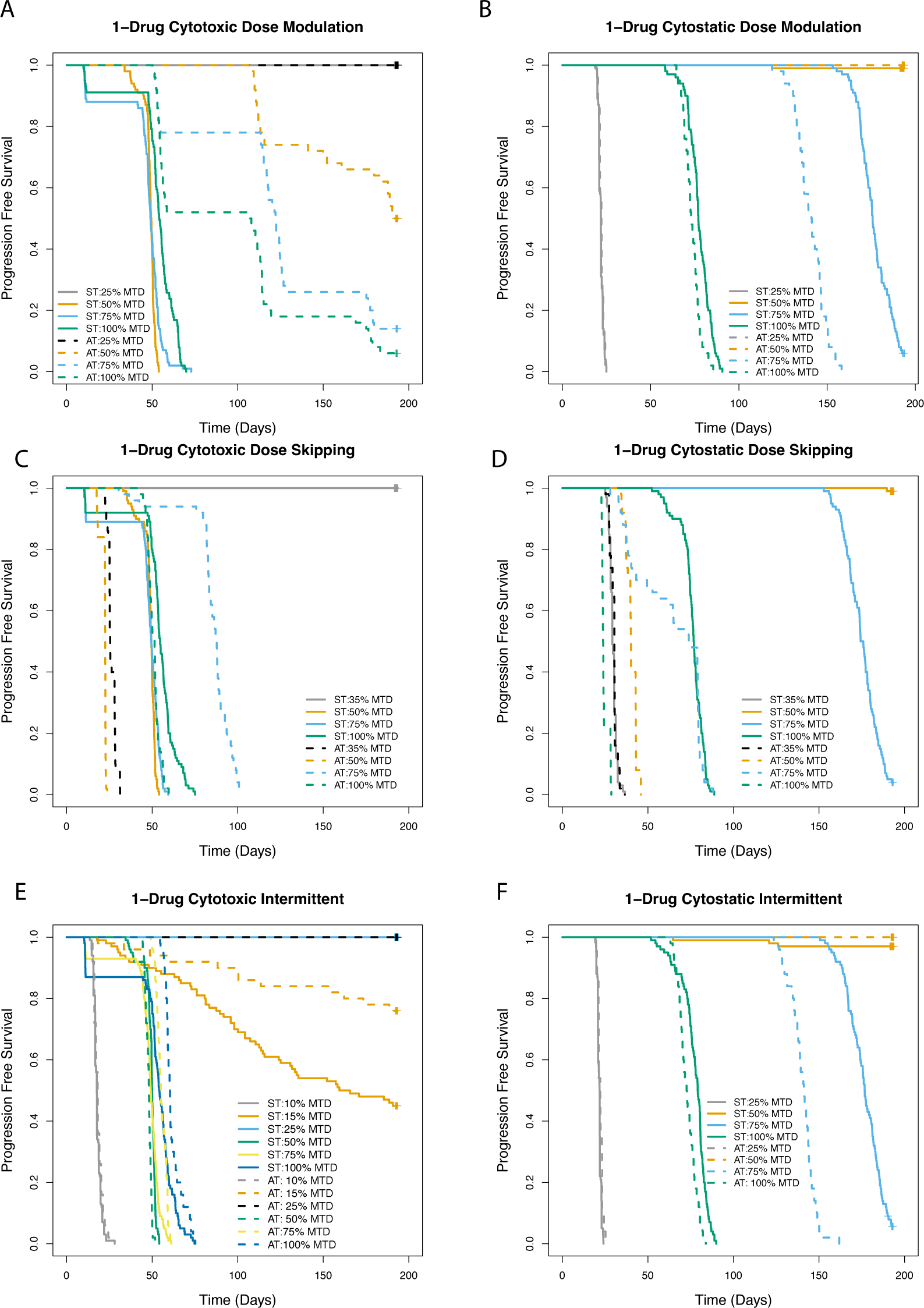
Effect of administering treatment at a range of different drug dosages for adaptive therapy using a single cytotoxic or a single cytostatic drug. Survival outcome comparing treatment as per the dose modulation protocol relative to ST, starting and capping dosing at 25%, 50%, 75%, or 100% of MTD, for treatment using a single cytotoxic drug (Fig. 9A), or a single cytostatic drug (Fig. 9B). Survival outcome for treatment as per dose-skipping protocol administered at 35%, 50%, 75%, or 100% of MTD relative to standard treatment using either a single cytotoxic drug (Fig. 9C), or a single cytostatic drug (Fig. 9D). Survival outcome for treatment as per the intermittent protocol administered at 10%, 15%, 25%, 50%, 75%, or 100% of MTD relative to ST for treatment using a single cytotoxic (Fig. 9E), or a single cytostatic (Fig. 9F) drug.

**Table 8:** Effect of administering treatment at a range of different drug dosages for adaptive therapy using a single cytotoxic or a single cytostatic drug

| Experimental Condition | Comparison Condition | Hazard Ratio | 95% CI | p-value |
| --- | --- | --- | --- | --- |
| **Dose Modulation (cytotoxic)** |  |  |  |  |
| 25% MTD | Standard Treatment at 25% MTD |  |  | Not Significant |
| 50% MTD | Standard Treatment at 50% MTD | ~0 |  |  |
| 75% MTD | Standard Treatment at 75% MTD | 0.06 | 0.03-0.12 | <0.001 |
| 100% MTD | Standard treatment at 100% MTD | 0.23 | 0.15-0.37 | <0.001 |
| **Dose Skipping (cytotoxic)** |  |  |  |  |
| Standard Treatment at 35% MTD | 35% MTD | ~0 |  |  |
| Standard Treatment at 50% MTD | 50% MTD | ~0 |  |  |
| 75% MTD | Standard Treatment at 75% MTD | 0.014 | 0.004-0.048 | <0.001 |
| 100% MTD | Standard treatment at 100% MTD | 2.9 | 2.0-4.3 | <0.001 |
| **Intermittent (cytotoxic)** |  |  |  |  |
| 10% MTD | Standard Treatment at 10% MTD |  |  | Not Significant |
| 15% MTD | Standard Treatment at 15% MTD | 0.34 | 0.18-0.64 | <0.001 |
| 25% MTD | Standard Treatment at 25% MTD |  |  | Not Significant |
| 50% MTD | Standard treatment at 50% MTD | 3.3 | 2.3-4.9 | <0.001 |
| 75% MTD | Standard treatment at 75% MTD | 0.33 | 0.23-0.47 | <0.001 |
| 100% MTD | Standard treatment at 100% MTD | 0.48 | 0.34-0.68 | <0.001 |
| **Dose Modulation (cytostatic)** |  |  |  |  |
| 25% MTD | Standard Treatment at 25% MTD |  |  | Not Significant |
| 50% MTD | Standard Treatment at 50% MTD |  |  | Not Significant |
| 75% MTD | Standard Treatment at 75% MTD | 248.8 | 56.4-1096 | <0.001 |
| 100% MTD | Standard treatment at 100% MTD | 2.4 | 1.7-3.4 | <0.001 |
| **Dose Skipping (cytostatic)** |  |  |  |  |
| 35% MTD | Standard Treatment at 35% MTD |  |  | Not Significant |
| Standard Treatment at 50% MTD | 50% MTD | ~0 |  |  |
| Standard Treatment at 75% MTD | 75% MTD | ~0 |  |  |
| Standard treatment at 100% MTD | 100% MTD | ~0 |  |  |
| **Intermittent (cytostatic)** |  |  |  |  |
| 25% MTD | Standard Treatment at 25% MTD | 0.53 | 0.37-0.77 | <0.001 |
| 50% MTD | Standard treatment at 50% MTD | ~0 |  |  |
| 75% MTD | Standard treatment at 75% MTD | 86.3 | 36.5-204 | <0.001 |
| 100% MTD | Standard treatment at 100% MTD | 2.7 | 1.9-3.9 | <0.001 |

For the dose-skipping protocol, we tested different drug dosage levels for administering the drugs, that is, 35%, 50%, 75%, or 100% of MTD. We observed an increase in TTP relative to standard treatment only for drug dosage level at 75% of MTD, but no increase in TTP relative to standard treatment for the other drug dosage levels tested here, that is 35%, 50%, or 100% (Fig. 9C, Table 8). For treatment using a single cytostatic drug, none of the drug dosage levels tested resulted in increase in TTP relative to standard treatment (Fig. 9D).

For the intermittent protocol, we tested different drug dosage levels for administering the drugs, that is, 10%, 15%, 25%, 50%, 75%, or 100% of MTD for cytotoxic drugs and 25%, 50%, 75%, or 100% of MTD for cytostatic drugs. We observed an increase in TTP relative to standard treatment only for drug dosage levels of 15%, 75%, and 100% of MTD (Fig. 9E, Table 8). For treatment using a single cytostatic drug, as per the intermittent protocol, only treatment with a drug dosage level at 50% of MTD resulted in an increase in TTP relative to standard treatment (Fig. 9F, Table 8).

### Median TTP versus Average Drug Dose

We plotted log10 median TTP versus average drug dose including all data points except for which a median TTP is not available (since less than 50% of the test subjects has progressed) and fitted the curve to a quadratic plot. For treatment using a single cytotoxic drug (Fig. 10A), median R-squared value is 0.1593 and adjusted R-squared is 0.1398. For treatment using a single cytostatic drug (Fig. 10B), median R-squared value is 0.6667 and adjusted R-squared value is 0.6592.

**Figure 10:**
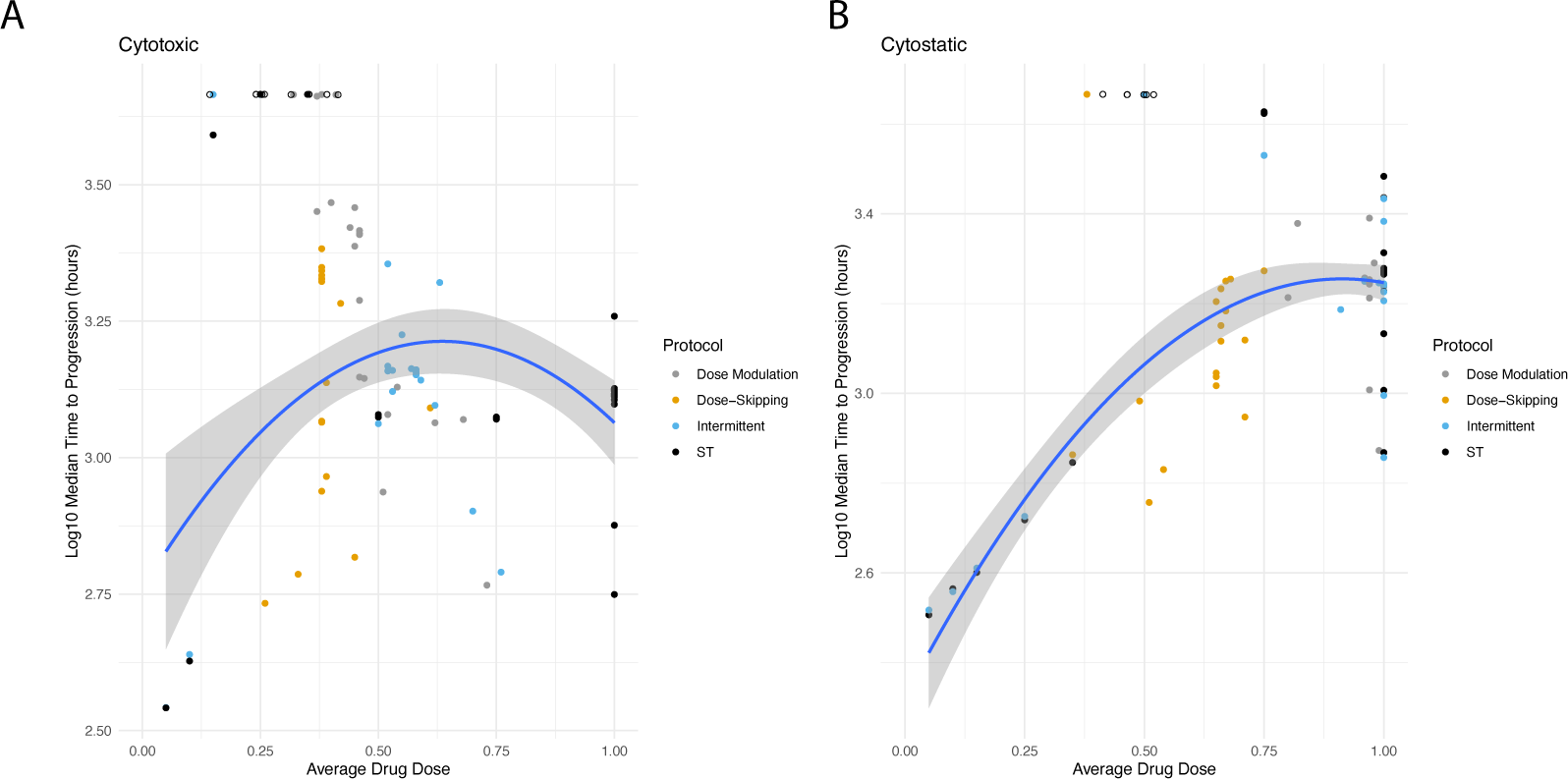
Summarizing the relationship between drug dose and time to progression for adaptive therapy using a single cytotoxic or a single cytostatic drug. In each panel, the average amount of drug used per timestep between the start of therapy and the time of progression is plotted on the X-axis, and the median time to progression for that protocol under those parameter values are plotted on the Y-axis, for treatment using either a single cytotoxic drug (A) or a single cytostatic drug (B). The points are colored based on the specific protocol. Open circles indicate data points that are censored as less than 50% of test-subjects have progressed, and are not included in the calculation. A quadratic fit to the curve along with the confidence intervals have been indicated in the figure panels. Each point represents a specific Kaplan-Meier survival curve for a given set of parameter values.

## Discussion

Design of adaptive therapy protocols for cancer treatment is challenging as, unlike standard treatment at maximum tolerated dose, where the same drug dosage is administered every treatment cycle, adaptive therapy treatment protocols typically involve multiple parameters, in order to account for the change in tumor burden and drug levels for every treatment cycle. Further, different drugs could differ in their mode of action, which could affect treatment outcome in important ways. Most of the literature involving mathematical modeling or agent-based simulations implicitly assume a cytotoxic mode of action. In this article, we sought out to investigate three different adaptive therapy protocols, namely, dose modulation, dose-skipping, and intermittent, and standard treatment at maximum tolerated dose, using either a single cytotoxic, or a single cytostatic drug, under a variety of different settings on cell kinetics, or treatment dynamics, with the goal of finding the optimum protocol for each setting, on a case-by-case basis. While cytotoxic drugs kill cells as a function of drug concentration, cytostatic drugs, as a function of drug concentration, prevents cells from dividing.

We observed all three adaptive therapy protocols tested here, that is, dose modulation, dose-skipping, and intermittent increased TTP relative to standard treatment, when using a single cytotoxic drug, but none of the protocols increased TTP relative to standard treatment when using a single cytostatic drug, under the default values of parameter settings. In line with the preclinical adaptive therapy experiments conducted in mice with breast cancer ^6^, which involved dosing with the drug paclitaxel, we see dose modulation outperforming dose-skipping using a single cytotoxic drug under the default parameter settings (HR=0.48, CI: 0.29-0.79, *p*<0.01). We did not observe increase in TTP relative to standard treatment for the intermittent protocol treatment using a single cytostatic drug, and thus our results do not agree with the prostate cancer clinical trials ^7^ conducted using the drug abiraterone, where the experimenters observed increase in TTP relative to a contemporaneous cohort of patients using the intermittent protocol. But our parameters are not calibrated to the individual prostate cancer patients ^7–9, 14^, and our default values of parameters may not represent the dynamics of that system. Furthermore, hormone therapy in prostate cancer appears to shrink the tumor and so may be having both a cytostatic and a cytotoxic effect.

Our results indicate fitness cost incurred by resistant cells, as manifested in longer doubling times relative to sensitive cells in the absence of the drug, is necessary, and higher fitness cost leads to improved survival outcome in most cases when using a single cytotoxic drug. However, when using a single cytostatic drug, despite a high fitness cost, none of the treatment protocols improve treatment outcome relative to standard treatment, with two exceptions, where we see dose-skipping outperforming standard treatment under 1.7-fold, or 2.5-fold fitness cost albeit with little effect size. Our results here are consistent to our findings in an earlier publication ^17^, where we observed a fitness cost of resistance is required for the multi-drug adaptive therapy protocols to work significantly better than standard treatment and that higher fitness cost translated to improved survival outcome.

The replacement parameter is an indicator of cell competition, as it specifies the likelihood that a cell can replace its neighbor if there are no empty spaces in its immediate neighborhood when it tries to divide. We observe under conditions of 0% replacement, when using a single cytotoxic drug, no increase in TTP relative to the standard treatment, but we observe an increase in TTP relative to standard treatment under conditions of 50%, or 100% replacement, with the singular exception where dose-skipping works poorly under conditions of 100% replacement, the progression being driven by total tumor burden and not the resistant cells in this case. Also, our observation that increase in replacement translates to an improved survival outcome, agrees with our earlier findings ^17^ that adaptive therapy survival outcomes using multiple drugs improves under conditions of higher replacement rates. When using a single cytostatic drug, however, no increase in TTP relative to the standard treatment was observed for any of the adaptive therapy protocols tested here, regardless of the level of replacement. Stem cells are able to replace each other ^22, 23^. Cancer cell cannibalism (entosis) is a phenomenon by which one cell kills its neighboring cell ^24–26^. Because cancer cell replacement leads to cell death, cell replacement can be considered to be a special kind of cell turnover, opening up spaces for the sensitive cells to proliferate at the expense of resistant cells in the absence of the drug potentially leading to improved survival outcome with cytotoxic drugs under conditions of higher replacement rates.

The turnover parameter is another indicator of cell competition, and it measures the effect of cell death and divisions with identical doubling times for each cell type across the two scenarios, that is low, or high turnover. In stark contrast to our findings earlier when we varied the replacement or the fitness cost parameter, where treatment using a single cytostatic drug did not lead to an increase in TTP relative to standard treatment for any of the adaptive therapy protocols tested, despite increase in fitness cost, or replacement, we observe every adaptive therapy protocol was able to increase TTP relative to standard treatment under conditions of high turnover, both for treatment using a single cytotoxic, or a single cytostatic drug. We find the turnover parameter to be critical, as it serves as an important determinant to whether treatment outcome would be better with an adaptive therapy protocol versus standard treatment. As such, markers for cell turnover would be paramount to assess the potential efficacy of adaptive therapy protocols. Strobl et al. has shown that turnover amplifies the effect of fitness cost of resistance ^18, 27^ by extending TTP. In our earlier publication ^17^ we found high turnover to either increase TTP relative to standard treatment for DM cocktail tandem, or insignificant with respect to low turnover for the two ping-pong protocols tried there (DM Ping-Pong on Progression, DM Ping-Pong Every Cycle). It could be that selection for the doubly resistant cells was stronger when both drugs were used simultaneously (DM Cocktail Tandem) and less when either of the drugs were used at a time, and thus the effect of turnover wasn’t noticeable as it maxed out for the two ping-pong protocols explored there. Also, there were 4 different cell types in our earlier model, in contrast to the 2 cell types here.

The dose modulation protocols have two primary parameters: Delta Tumor, which is the amount the tumor burden must change in order to trigger a change of drug dose and, Delta Dose, which is the amount by which the drug dose is changed. Similar to our findings in the 2-drug paper, we observe all of the adaptive therapy protocols work best when using a relatively low value of delta tumor versus a high value. As such, we observe the general trend that survival outcomes decrease in the following order for the delta tumor parameter: 5%, 10%, 20%, and 40%. This observation can also be extended to treatment using a single cytostatic drug, as dose modulation protocol using a single cytostatic drug improves survival outcome relative to standard treatment only when a delta dose value=5% was used, and not under delta dose=10%,20%, or 40%. For the delta dose parameter, however, we observe improved survival outcome when a higher delta dose value is used, with the best survival outcome at delta dose=75%, for both treatment using a single cytotoxic, or a single cytostatic drug. These results are in line with our earlier findings ^17^, where we noted poor survival outcomes when using a too conservative value for the delta dose parameter. In another study ^2^, a single drug adaptive therapy regimen with the dose modulation protocol with Delta Tumor=10% and Delta Dose=50% was shown to work better than Delta Tumor=5% and Delta Dose=25%. Our modeling results suggest that dose modulation with Delta Tumor=5% and Delta Dose=75% would work best for treatment using a single cytotoxic, or a single cytostatic drug.

An open question in the field of adaptive therapy is when to withhold treatment, or in other words to back off on the drug, when it is indeed feasible to do so. For intermittent treatment protocols, a key question is at what tumor burden should the treatment be stopped when the tumor is shrinking, in order that the tumor may be allowed to climb back up to the baseline value at which treatment was initiated previously. For treatment using the dose modulation protocol, we have to decide whether to withhold treatment when the tumor is shrinking or, to continue adjusting the drug dosages. For treatment as per the intermittent protocol, for both treatment using a single cytotoxic, or a single cytostatic drug, the tumor burden relative to the baseline at which we stop dosing has no significant effect. For treatment using the dose modulation protocol, however, we observe withholding treatment when the tumor shrinks by 20% to be far better than waiting to withhold treatment until the tumor shrinks by 50%. For treatment using a single cytostatic drug, however, these effects were not significant. These results agree with our findings ^17^ earlier, where we found incorporating frequent treatment vacations works best for DM Cocktail Tandem. Interestingly, it has been shown ^17, 18^ that treatment vacations would provide a benefit only under conditions of strong intra-tumoral competition. However, in that publication an ordinary differential equation was used and spatial effects were not studied.

We also explored what is an optimum drug dosage level to administer at therapy initiation (for the dose modulation protocol), or at each treatment cycle (for intermittent and dose-skipping). In general, a drug dosage level at 50%, or 75% of the MTD works well for both treatment using a single cytotoxic, or a single cytostatic drug. In addition, fixed dosing with a cytostatic drug at 50% of MTD worked almost all the time, with only a few failures. We also found a variety of protocols that were able to control the tumors indefinitely:

1. Cytotoxic dose modulation starting and capped at 25% MTD (with DeltaTumor 10%, DeltaDose 50%, and stopping treatment at 50% of the initial tumor burden)
2. Cytotoxic intermittent using a fixed dose at 25% of MTD (and stopping dosing at 50% of the initial tumor burden, and restarting when it recovers to 100% of the initial tumor burden)
3. Cytotoxic fixed dosing at 25% of MTD (regardless of how the tumor responds)
4. Cytostatic intermittent with a fixed dose at 50% of MTD
5. Cytostatic dose modulation starting and capped at 50% of MTD

To some extent, standard treatment protocols at doses less than 100% of the maximum tolerated dose (such as standard treatment at 10%, 15%, 25%, 35%, 50%, or 75% of MTD) are more along the lines of metronomic therapy albeit that the treatment frequency remains the same. The effects of cancer treatment using metronomic scheduling has been studied ^28, 29^. Our observation here that standard treatment could lead to improved survival outcome provided a fraction of maximum tolerated dose is used has important implications as it suggests a more personalized patient-centered approach to treatment has the potential to work significantly better in clinical settings. Furthermore, because we did not consider toxicity in our models, the benefit observed using standard treatment at low drug dosages can be reasonably expected to have been underestimated and should work even better in clinical and experimental settings.

One general principle that emerged from these simulations is that there is a sort of Goldilocks level of drug exposure. If too much drug is used, there is strong selection for resistance, and we lose control of the tumor due to the resistant clones growing out. However, if too little drug is used, we cannot keep control of the sensitive cells and the tumor grows out of control. When we analyzed the relationship between the average amount of drug used per unit time and the time to progression, we found a significant unimodal relationship, fitting this Goldilocks principle ^30^. The R-squared values on those regressions are consistent with the fact that there are many determinants of time to progression, but the average amount of drug exposure per unit of time is clearly a significant factor. A Goldilocks level of drug dosage has been observed to be optimal in tyrosine kinase inhibitors (pazopanib, which is a VEGF receptor TKI) in a clinical setting in patients with advanced renal cell carcinoma ^31^

It is not clear if drugs that are putatively cytostatic are acting as truly cytostatic drugs. For e.g., targeted therapies that affect growth factor receptors and hormone therapies for breast and prostate cancer have been shown to actually shrink the tumor. The cell killing effect of cytostatic drugs is actually due to oncogene or hormone addiction. And thus, when you take off the drug, the cells die due to the cell killing effect. Look up some oncogene and hormone addiction papers.

Our work has several limitations. We generally don’t have the technology to accurately measure total tumor burden changes of 5%. We also often do not have cost effective ways to carry out those measures frequently. However, we are essentially trying to control a complex system, and lag times between changes in the system and control responses often lead to loss of control. In the future, we will be exploring 2-drug cytostatic adaptive therapy protocols.

## Conclusions

Dose modulation, dose-skipping, as well as intermittent treatment protocols work well under a wide range of parameter settings when treating using a single cytotoxic drug. In contrast, there seems to be only a handful of parameter settings that improves survival outcome when using a single cytostatic drug. In general, adaptive therapy, using either a single cytotoxic, or a single cytostatic drug, works best under conditions of high competition among the cell types, such as higher fitness cost, high levels of replacement, or high turnover. Our results suggest assaying for the amount of turnover in the cancer would be helpful for determining the likely efficacy of adaptive therapy. In general, there seems to be an intermediate level of drug we can use, which maximizes TTP, as too little leads to progression of sensitive cells and too much leads to progression of resistant cells. In fact, we found that even a constant dosing of an intermediate drug level can provide long term control even without using adaptive therapy. Our results suggest that cancer therapy could be significantly improved by the development of sensitive and accurate measures of tumor burden, that can be used frequently to track tumor response to therapy. We should note that adaptive therapy is most appropriate when the presence or emergence of therapeutic resistance is likely and cure is unattainable. If successful, adaptive therapy holds the promise of changing cancer from an acute lethal disease into a chronic disease that does not kill us.

## Notes

### Competing Interest Statement

The authors have declared no competing interest.

